# Pancreatic ductal adenocarcinoma cells employ integrin α6β4 to form hemidesmosomes and regulate cell proliferation

**DOI:** 10.1101/2021.08.19.456969

**Authors:** Jonathan D. Humphries, Junzhe Zha, Jessica Burns, Janet A. Askari, Christopher R. Below, Megan R. Chastney, Matthew C. Jones, Aleksandr Mironov, David Knight, Derek A. O’Reilly, Mark J. Dunne, David R. Garrod, Claus Jorgensen, Martin J. Humphries

## Abstract

Pancreatic ductal adenocarcinoma (PDAC) has a dismal prognosis due to its aggressive progression, late detection and lack of druggable driver mutations, which often combine to result in unsuitability for surgical intervention. Together with activating mutations of the small GTPase KRas, which are found in over 90% of PDAC tumours, a contributory factor for PDAC tumour progression is formation of a rigid extracellular matrix (ECM) and associated desmoplasia. This response leads to aberrant integrin signalling, and accelerated proliferation and invasion. To identify the integrin adhesion systems that operate in PDAC, we analysed a range of pancreatic ductal epithelial cell models using 2D, 3D and organoid culture systems. Proteomic analysis of isolated integrin receptor complexes from human pancreatic ductal epithelial (HPDE) cells predominantly identified integrin α6β4 and hemidesmosome components, rather than classical focal adhesion components. Electron microscopy, together with immunofluorescence, confirmed the formation of hemidesmosomes by HPDE cells, both in 2D and 3D culture systems. Similar results were obtained for the human PDAC cell line, SUIT-2. Analysis of HPDE cell secreted proteins and cell-derived matrices (CDM) demonstrated that HPDE cells secrete a range of laminin subunits and form a hemidesmosome-specific, laminin 332-enriched ECM. Expression of mutant KRas (G12V) did not affect hemidesmosome composition or formation by HPDE cells. Cell-ECM contacts formed by mouse and human PDAC organoids were also assessed by electron microscopy. Organoids generated from both the PDAC KPC mouse model and human patient-derived PDAC tissue displayed features of acinar-ductal cell polarity, and hemidesmosomes were visible proximal to prominent basement membranes. Furthermore, electron microscopy identified hemidesmosomes in normal human pancreas. Depletion of integrin β4 using siRNA reduced cell proliferation in both SUIT-2 and HPDE cells, reduced the number of SUIT-2 cells in S-phase, and induced G1 cell cycle arrest, indicating a requirement for α6β4-mediated adhesion for cell cycle progression and growth. Taken together, these data suggest that laminin-binding adhesion mechanisms in general, and hemidesmosome-mediated adhesion in particular, may be under-appreciated in the context of PDAC.

Proteomic data are available via ProteomeXchange with the identifiers PXD027803, PXD027823 and PXD027827.

## Introduction

Pancreatic ductal adenocarcinoma (PDAC) is one of the five most common causes of cancer mortality in developed countries and exhibits one of the worst clinical outcomes [1,2]. A prominent feature of PDAC is an extensive desmoplastic reaction that makes a multifactorial contribution to tumour progression and disease lethality. In PDAC, where the stroma on average constitutes 80% of total tumour volume, desmoplasia is exaggerated compared to other carcinomas[3]. As a consequence, the pathologically remodelled and rigid ECM in PDAC desmoplasia leads to aberrant integrin signalling, resulting in accelerated proliferation and invasion [4–8].

Integrins are a family of cell surface receptors that mediate adhesion to the ECM and form connections to the cytoskeleton [9,10]. In addition to providing a structural connection, integrins act as bidirectional signalling hubs relaying biochemical and biomechanical signalling pathways to regulate cell adhesion and modulate a range of phenotypic outputs [11]. Integrin activation and/or ligand binding lead to the formation of plasma membrane-localised protein complexes, termed integrin adhesion complexes (IAC) [10,12,13]. IACs function as mechanosensitive molecular clutches that transmit forces between the ECM and cytoskeleton [14,15]. Data from both literature curation [16,17] and proteomic analysis [18–25] demonstrate that a small number of proteins establish the framework of the IAC adhesome, and a larger cohort of more transient proteins tune its function to intra- and extracellular stimuli [26,27]. In this way, it is hypothesised that individual components of IACs act in a cooperative manner to provide coordinated functional adhesion signalling outputs [12].

Based on our understanding of the integrin adhesion-dependent control of cell fate, the generation of a rigid ECM would be likely to alter proliferation, invasion and differentiation [28], and there is evidence for this in PDAC [29,30]. In addition, integrins and the ECM are known to contribute to the hallmarks of cancer [31,32]. There is also growing evidence for a mechanistic coupling of integrin function with the cell cycle to govern cell proliferation [33–35], and inhibition of the IAC component focal adhesion kinase limits tumour progression in the KPC mouse model of human PDAC [36]. Integrins and IACs are therefore considered important regulators of the pathological development of cancer and provide opportunities for therapeutic intervention [7,8,31,37,38]. Elucidating the mechanisms employed by PDAC cells to interact with the desmoplastic ECM would therefore be important in the quest to improve patient outcomes.

## Results

### HPDE cell adhesion receptor complexes are dominated by integrin α6β4 and hemidesmosome components

The most prevalent mutations in PDAC, observed in over 90% of all cases are in KRas, a small GTPase implicated in a wide range of signalling pathways [39]. As a first step to understanding the adhesome composition of PDAC, we characterised the H6c7 normal human pancreatic ductal epithelial (HPDE) cell model. This permitted a comparison of matched wild-type (control) and mutant human KRas expressing (KRas G12V) HPDE cell lines [40,41]. Increased Ras activity was confirmed in KRas G12V HPDE cells compared to control cells (Fig 1A). Cells were grown in monolayer culture for seven days to enable assembly of cell-derived ECM, and then IAC isolation was carried out for both cell lines and the samples subjected to mass spectrometry (MS)-based proteomic analysis [26,42]. Although we employed our standard IAC isolation protocol [42], the crosslinking step, using DTBP, was not required for HPDE cells.

**Figure 1:**
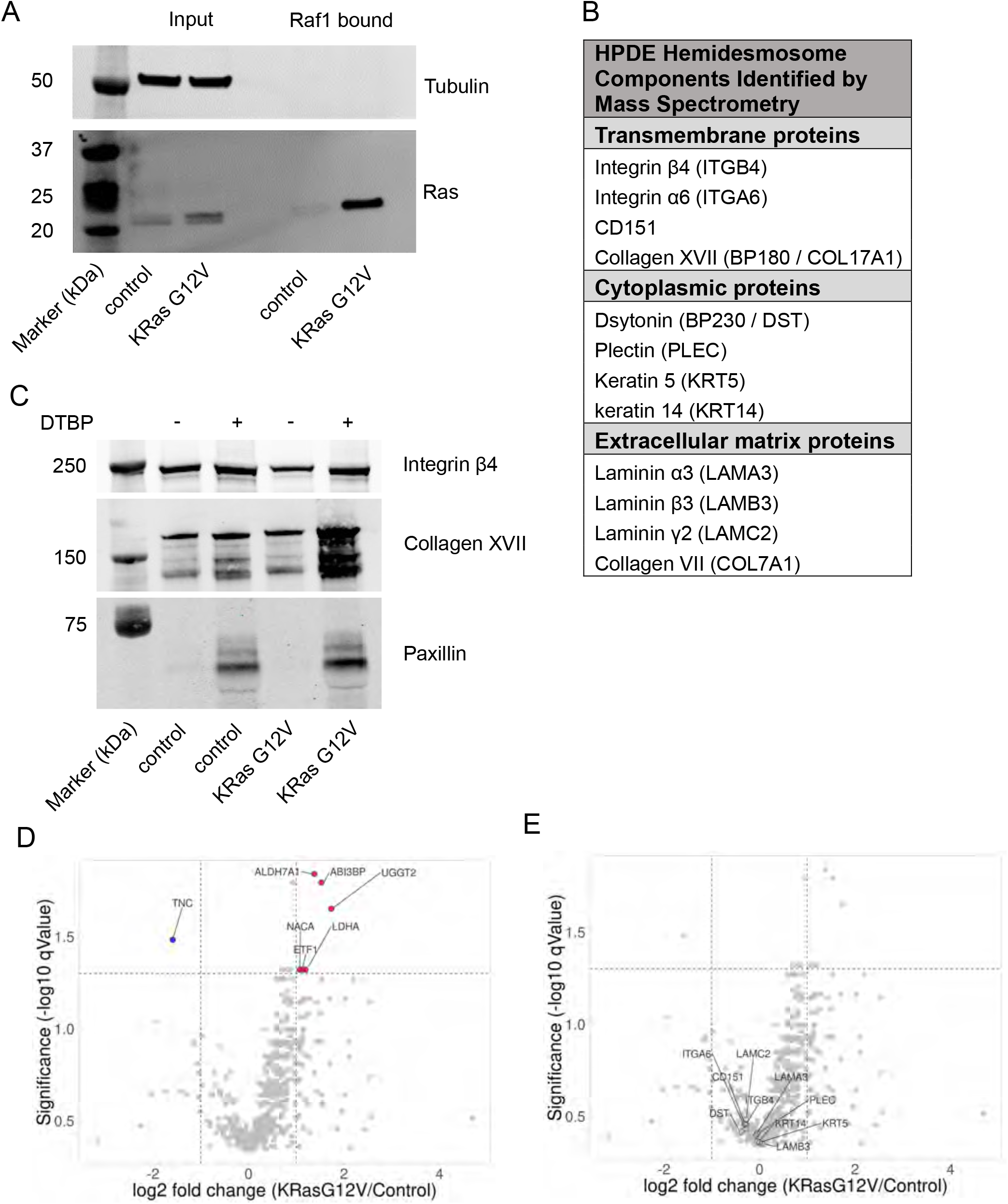
HPDE cells form α6β4-based adhesion complexes containing hemidesmosome components. (A) Ras activity was determined in wild-type (control) and mutant KRas expressing (KRas G12V) HPDE cells using the GST-Raf1-RBD effector pulldown assay. Active Ras was detected in the Raf1 bound fraction and tubulin was used to demonstrate specificity. (B) IACs were isolated from control and KRas G12V-expressing HPDE cells after 7 days in culture and subjected to MS analysis. Hemidesmosome components identified from HPDE IAC isolations by MS are listed. (C) Western blotting confirmed the identification of integrin β4 and collagen XVII, but not paxillin, from HPDE IAC isolations. The use of the crosslinking reagent (DTBP) is indicated above. (D) and (E) MS-based abundance ratios (KRasG12V/control) and qValues calculated by Progenesis QI (as described in Methods) for proteins detected in HPDE IAC isolations and displayed as volcano plots (Supp. Table 1). Panel (D) displays proteins with significantly altered abundance profiles (red = increased in KRas G12V and blue = decreased in KRas G12V IACs). Panel (E) displays selected hemidesmosomal proteins, demonstrating that they do not significantly change between KRas G12V and control HPDE IACs. For western blots, the sizes of molecular weight markers are indicated to the left of images.

MS analysis detected 576 proteins from all conditions (Supp Table 1). Comparison to the *in silico* literature-based integrin adhesome [16,17,43] revealed 29 adhesome proteins (12.5% of 232), with 16 intrinsic and 13 associated components. This is consistent with the coverage of the adhesome achieved from other integrin adhesome complex isolations (range, 9.1–32.3% [26]); however, in contrast to data from other cell types, the only integrins robustly identified in the HPDE datasets were the α6 and β4 subunits, suggesting that H6c7 HPDE cells unexpectedly employ integrin α6β4 to adhere to the ECM. Consistent with the detection of α6β4, further analysis of the HPDE datasets revealed that the some of the most abundant proteins detected in H6c7 IACs were components of hemidesmosomes and the associated cytokeratin-based cytoskeletal network (fig 1B[44]). This was supported by Gene Ontology (GO) analysis, which demonstrated an enrichment for terms such as hemidesmosome (28.5-fold enrichment; GO:0030056) and hemidesmosome assembly (30.4-fold enrichment; GO:0031581). In contrast, relatively few proteins (13/70 = 18.6%) were detected from the consensus adhesome (integrin adhesion complex components commonly identified from cells interacting with fibronectin [26]). 350 of the 576 identified HPDE IAC components (60.7%) were members of the meta-adhesome (i.e. enriched in seven published IAC datasets [26]). These analyses suggest that the HPDE adhesome is a highly cell type-dependent adhesome variant, possibly reflecting differences between epithelial HPDE cells, and mesenchymal cells used to inform eh meta-adhesome.

Western blotting was employed to support the data from isolation of HPDE IACs. Integrin β4 and collagen XVII were detected in HPDE IACs without the requirement for DTBP crosslinker, but not the consensus adhesome component paxillin (fig 1C), highlighting the specificity of the isolated HPDE IAC adhesome. Moreover, relatively few of the identified IAC proteins displayed altered abundances upon expression of mutant KRas G12V (fig 1D), and the abundance of the hemidesmosome components was not significantly changed (fig 1E). These data indicate that normal HPDE cells are likely to form hemidesmosomes in culture, and the abundance of hemidesmosome components at ventral membrane sites is not altered by expression of mutagenic KRas G12V.

### HPDE cells secrete and form a laminin-rich matrix

From the GO analysis of HPDE IACs, it was noted that the identified ECM components (87 matrisome components; 8.19% matrisome coverage) [43,45,46] were enriched for basement membrane components (11.5-fold enrichment; GO:0005604∼basement membrane) including the integrin α6β4-binding laminins (laminin-332, laminin-511 and laminin-521), nidogen 1, collagen IV and perlecan.

To test the possibility that HPDE cells secrete and form a laminin-rich ECM, two additional MS-based proteomic approaches were used to identify the secreted ECM and CDM proteins (Supp Tables 2 and 3). The CDM analysis identified 701 proteins, and the secreted protein analysis identified 902 proteins. Both approaches achieved a similar coverage of the core matrisome; however, the secreted protein analysis identified more matrisome-associated proteins (fig 2A). Therefore, the combination of both approaches led to the identification of a more complete set of ECM proteins produced by HPDE cells. Gene Ontology (GO) analysis of both datasets supported the enrichment of ECM proteins including basement membrane proteins (Supp Table 4). These data demonstrate that H6c7 HPDE cells secrete and assemble laminin-rich basement membrane type ECM components in culture. In support of the identification of hemidesmosome components from HPDE cells, laminin-332 subunits were an abundant component of both the secreted protein and CDM datasets. The CDM analysis also robustly identified integrin α6β4 and other hemidesmosome components (collagens VII and XVII, and dystonin), confirming the localisation to HPDE cell ventral membranes by an alternative enrichment strategy. In agreement with the analysis of hemidesmosome components from isolated IACs, relatively few of the identified ECM proteins from secreted protein or CDM samples displayed altered abundances upon expression of mutant KRas G12V (fig 2B,C), and the abundance of all hemidesmosome-associated ECM components was not significantly changed (fig 2D,E).

**Figure 2:**
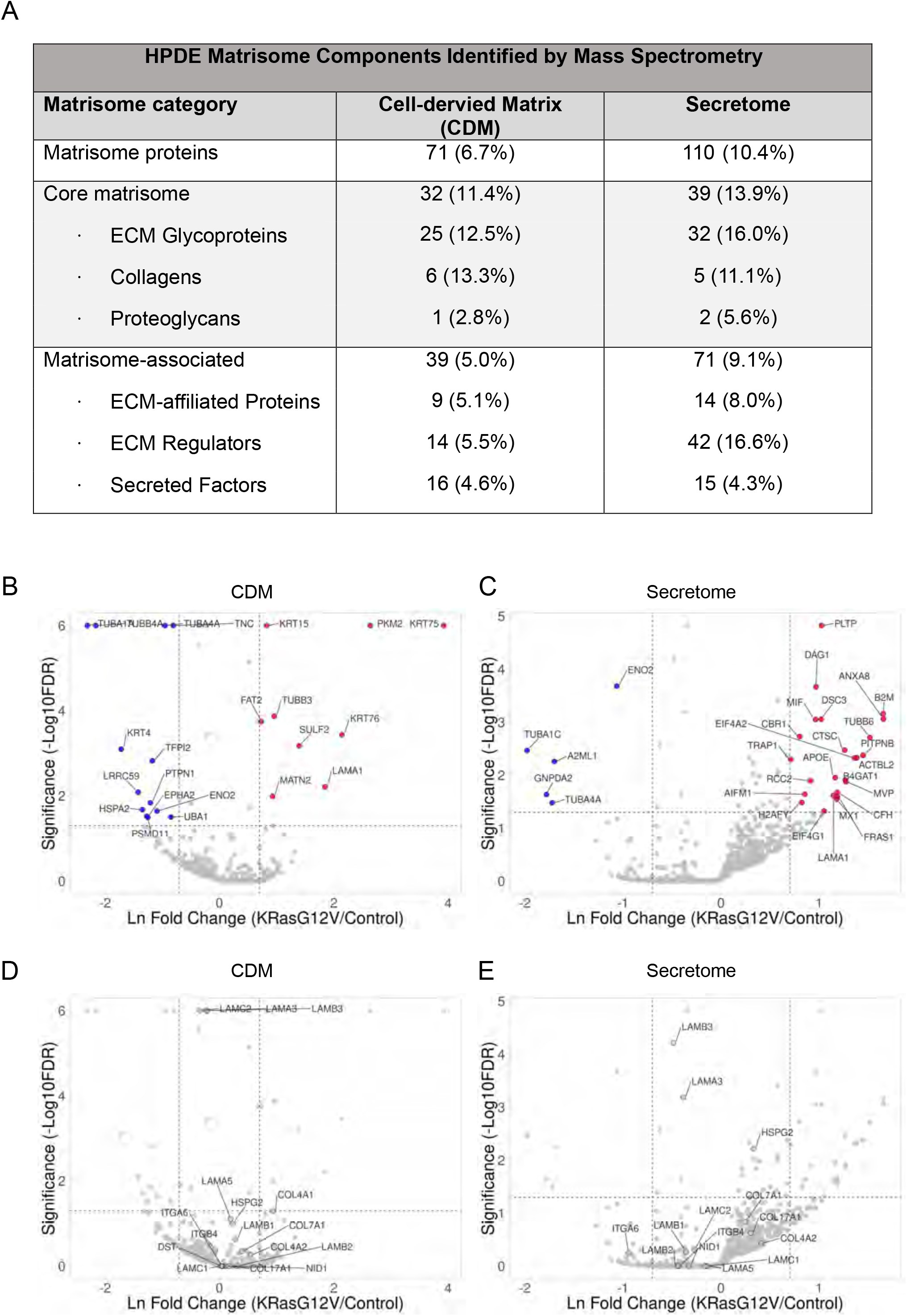
HPDE cells produce a laminin-based extracellular matrix. (A) CDMs and secreted proteins were isolated from control and KRas G12V-expressing HPDE cells after 7 days in culture and subjected to MS analysis. Numbers of proteins assigned to matrisome component categories are listed (see Supp. Tables 2 and 3). (B), (C), (D) and (E) MS-based abundance ratios (KRasG12V/control) and FDRs calculated by QSpec (as described in Methods) for proteins detected in HPDE CDM and secreted protein samples are displayed as volcano plots (Supp. Tables 2 and 3). Panels (B) and (C) display proteins with significantly altered abundance profiles (red = increased in KRas G12V and blue = decreased in KRas G12V) for CDMs and secreted proteins as indicated. Panels (D) and (E) displays selected basement membrane hemidesmosomal proteins, demonstrating that they do not significantly change between KRas G12V and control HPDE IACs.

### HPDE cells express α6β4 and form hemidesmosomes

Overall, the MS-based proteomic analysis of HPDE adhesion complexes and ECM suggested that hemidesmosome components act as the main adhesion machinery used by HPDE cells, and this was not altered by the expression of mutant KRas G12V. Next, we sought to verify the cell surface expression of integrin α6β4 and the formation of hemidesmosomes by flow cytometry, co-immunoprecipitation, immunofluorescence microscopy and electron microscopy.

Flow cytometry using a panel of antibodies directed against a range of integrin subunits and heterodimers revealed the cell surface expression of α2, α3, α5, α6, αV, β1 and β4, but not α1, α4, αVβ3 or αVβ5, in both control and KRas G12V-expressing HPDE cells. These results are consistent with the expression of integrin heterodimers acting as collagen receptors (α2β1), laminin receptors (α3β1, α6β1 and α6β4), and fibronectin receptors (α5β1 and αVβ1) in HPDE cells. Immunopreciptation using anti integrin α6, β1 and β4 antibodies confirmed the preferential association of α6 with the β4 subunit compared to β1 (Fig S1), which is consistent with the proteomic identification of α6β4 from HPDE cell adhesion complexes (Fig 1).

To test for the presence of integrin α6β4-based adhesion complexes in HPDE cells, immunofluorescence imaging was performed (Fig 3). The integrin β4 subunit was observed in a characteristic leopard skin pattern, which is classically associated with hemidesmosomes [47], for both control and KRas G12V HPDE cells (Fig 3A). To assess the localisation of other hemidesmosome components, control HPDE cells were stained using antibodies to collagen XVII (BP180), dystonin (BP230) and collagen VII (Fig 3B). These analyses showed similar subcellular localisations as integrin β4, which is consistent with their detection in isolated IACs by MS and the formation of hemidesmosomes in HPDE cells.

**Figure 3:**
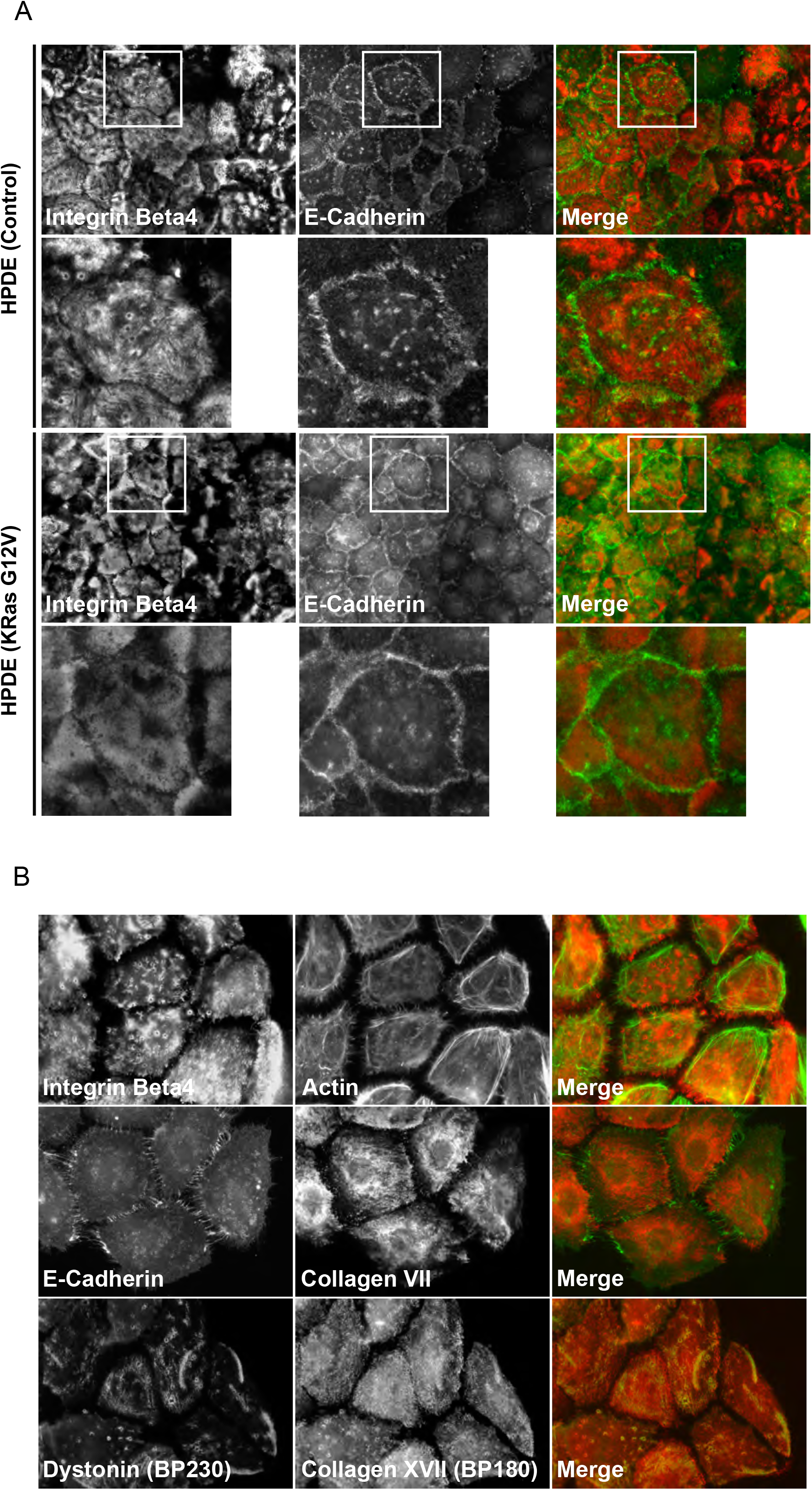
HPDE cells form characteristic hemidesmosome-type adhesion structures. (A) and (B) HPDE cells were cultured for up to 7 days on glass coverslips and immunofluorescence imaging performed. (A) Control and KRas G12V-expressing HPDE cells stained for integrin β4 displayed the same characteristic leopard skin pattern whilst E-cadherin stained cell-cell junctions. (B) Control HPDE cells were stained using antibodies directed against integrin β4, E-cadherin, collagen VII, dystonin (BP230) and collagen XVII (BP180).

The definitive demonstration of hemidesmosome formation is achieved by transmission electron microscopy (TEM), and the observation of the classical hemidesmosome ultrastructural organisation [44]. To this end, transverse sections of the HPDE-ECM interface were prepared and imaged by TEM (Fig 4). In general, HPDE cells displayed a morphology with a flat cell-ECM interface, microvilli on their dorsal surface and cell-cell contacts that were highly interdigitated. These analyses revealed abundant, electron-dense hemidesmosome structures at the ECM interface of HPDE cells that linked directly to prominent cytokeratin filaments. The hemidesmosome structures formed over a period of one to eight days, but were infrequently observed at earlier time points and increased in frequency and maturity from three to six days. Hemidesmosomes were observed in both the presence or absence of expression of mutant KRas G12V, and comprised all the classically defined zones, including the juxtamembrane cytoplasmic inner and outer plaques, the extracellular lamina lucida and lamina densa, along with anchoring fibrils and filaments that project into the ECM [44,48]. In summary, the combination of evidence provided by MS, immunofluorescence and electron microscopy demonstrates that HPDE cells in culture form type I hemidesmosomes, containing the full repertoire of components, and localise to the cell-ECM interface.

**Figure 4:**
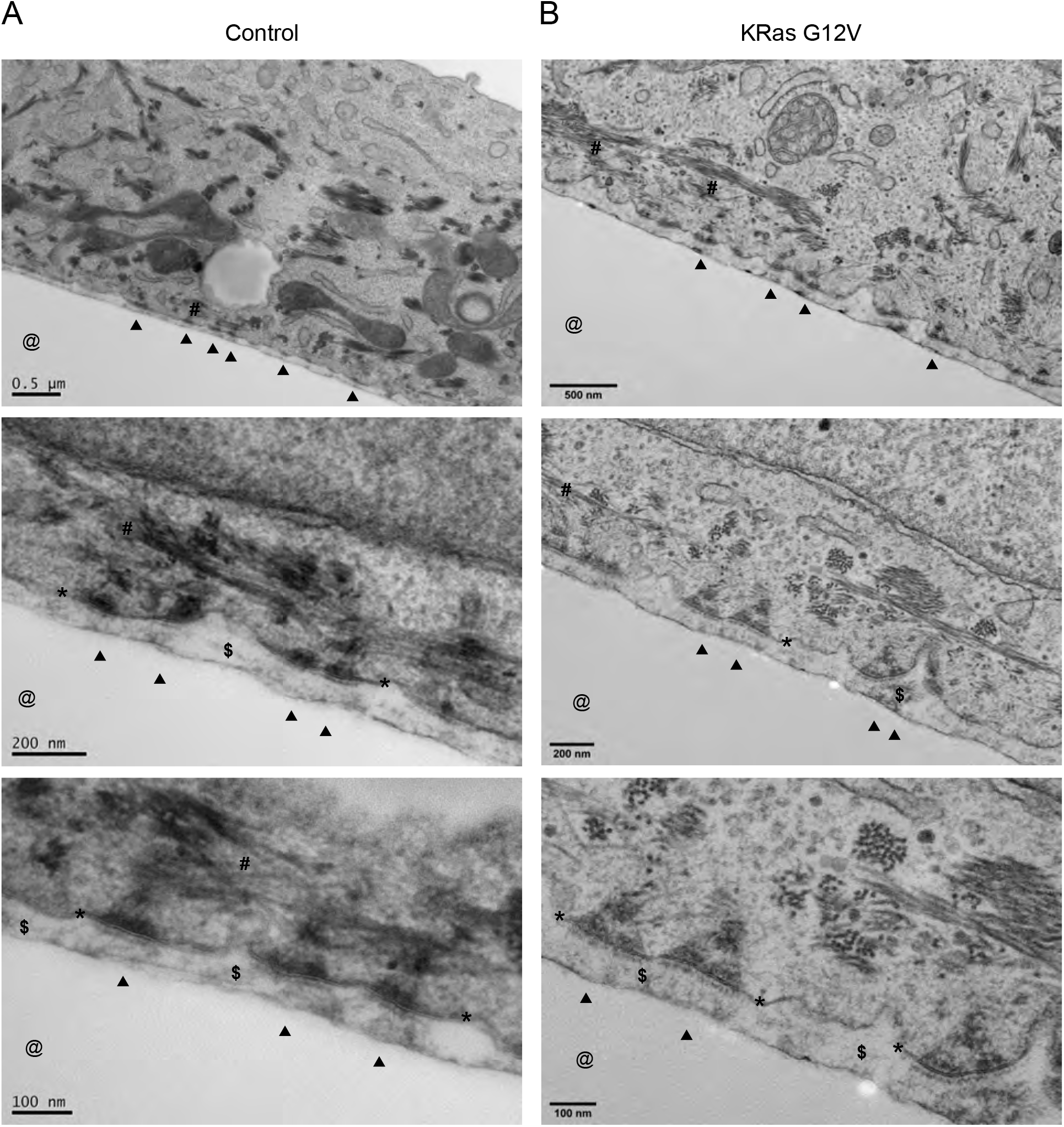
HPDE cells form hemidesmosomes in 2D culture. (A) Wild-type (control) and (B) mutant KRas-expressing (KRas G12V) HPDE cells were cultured on Aclar for up to 7 days. Transverse sections of the HPDE-ECM interface were prepared, imaged by TEM and a range of magnifications shown. Cells formed flattened basal surface with a thin layer of ECM (**$**) proximal to the area where the Aclar film (**@**) would have occupied. Arrowheads (▲) indicate the approximate position of some hemidesmosomes (indicated from the extracellular side) which are located at the plasma membrane (*) and link to cytoplasmic cytokeratin filaments (**#**). Images are orientated with the cell-ECM interface towards the bottom.

### HPDE cells form hemidesmosomes in 3D culture

3D cell culture systems offer a way to assess cell behaviour in environments that more closely mimic the spatial organisation and cell-ECM interactions *in vivo*, compared to 2D cell culture [49,50]. To assess the relevance of such culture conditions on the formation of hemidesmosomes in HPDE cells, H6c7 cells were grown in a 3D culture system that incorporated alginate and Matrigel, that had been used previously to investigate hemidesmosomes in mammary epithelial cells [51,52]. Both control and KRas G12V-expressing HPDE cells formed colonies of cells over 60 hours that expressed integrin β4, and the hemidesmosome component collagen XVII, that localised at the basal cell-ECM interface (Fig 5A) suggesting that HPDE cells form hemidesmosomes in 3D culture. To verify the presence of hemidesmosomes, HPDE cells grown in alginate/Matrigel 3D gels were processed and visualised by TEM. Electron-dense hemidesmosome structures were observed at cell-ECM interfaces of HPDE cells that were linked directly to cytokeratin filaments. Hemidesmosomes were observed in the presence or absence of expression of mutant KRas G12V (Fig 5B). These data demonstrate the formation of hemidesmosome structures by HPDE cells in 3D culture was not altered by the expression of mutant KRas G12V.

**Figure 5:**
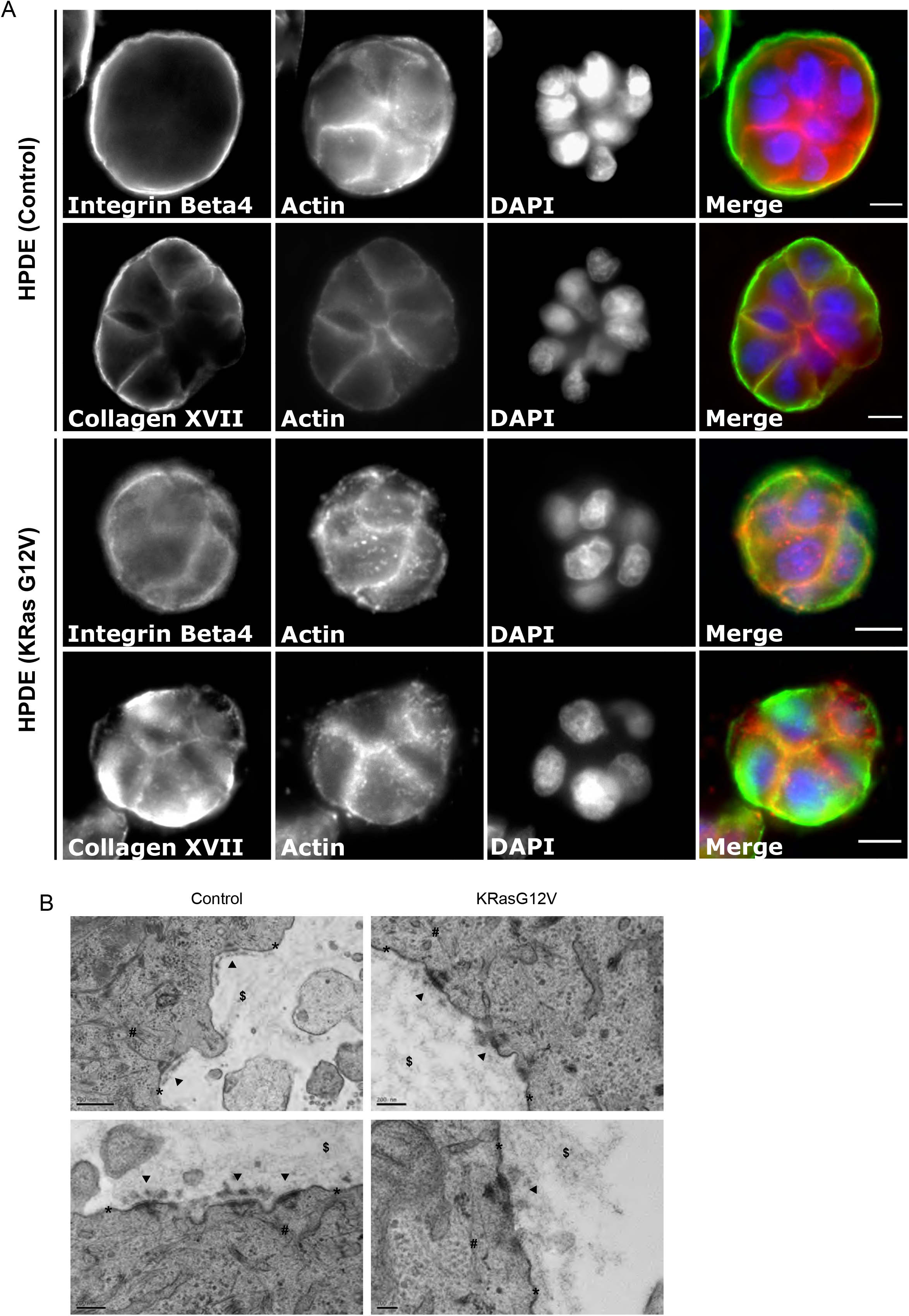
HPDE cells form hemidesmosomes in 3D culture. (A) Wild-type (control) and mutant KRas-expressing (KRas G12V) HPDE cells were grown in 3D culture (alginate / Matrigel) for 60 hours and immunofluorescence imaging performed. Control and KRas G12V-expressing HPDE cells stained for integrin β4 or collagen XVII. Scale bars represent 10 µm. (B) TEM was performed for wild-type (control) and mutant KRas-expressing (KRas G12V) HPDE cells after 60 hours in 3D culture (alginate / Matrigel). Arrowheads (▲) indicate the approximate position of some hemidesmosomes (indicated from the extracellular side) which are located at the plasma membrane (*), and often positioned proximal to a layer of basement membrane. The general position of the alginate / Matrigel ECM (**$**) along with cytoplasmic cytokeratin filaments (**#**).

### Murine and human PDAC organoids and normal human pancreas form hemidesmosomes

PDAC organoids have shown promise as they recapitulate the full spectrum of tumour development [53–55]. To develop further our understanding of PDAC-specific adhesion systems, the cell-ECM contacts used by mouse and human organoids were assessed by TEM. Organoids generated from both the KPC mouse model and human patient-derived PDAC tissue displayed keys features of acinar-ductal cell polarity with appropriately positioned luminal microvilli, tight junctions and adherens junctions (Fig 6A). Moreover, hemidesmosomes were detected in close proximity to prominent basement membranes in both KPC and human PDAC organoids (Fig 6B). Furthermore, the presence of hemidesmosomes in normal human pancreas was confirmed, at the ultrastructural level, positioned at the basal surface of ductal cells in close proximity to a prominent basement membrane (Fig 6C). The overall cellular organisation and ductal cell polarisation was indistinguishable between the human organoids and normal human pancreas.

**Figure 6:**
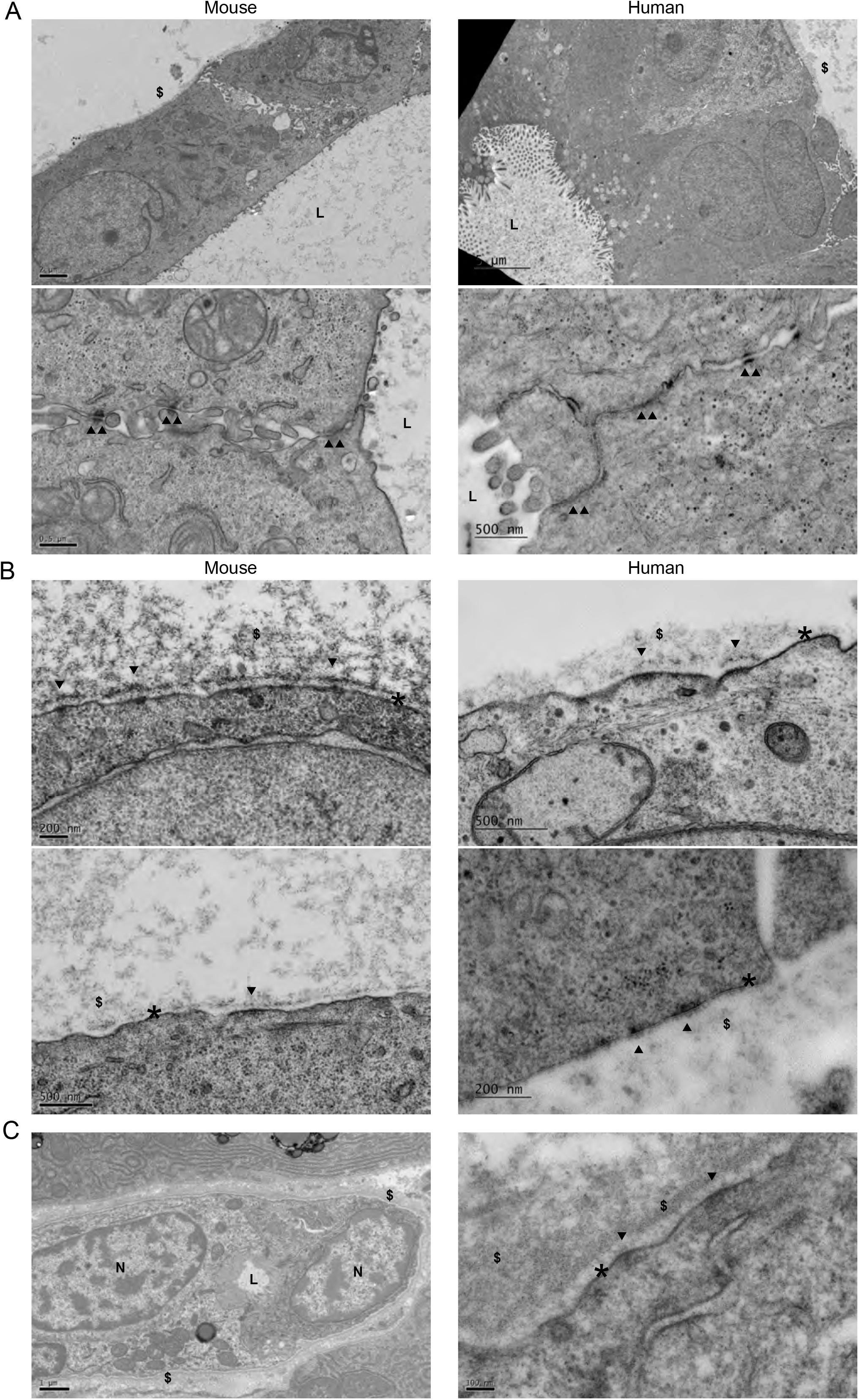
PDAC organoids and human pancreas form hemidesmosomes. (A) and (B) TEM was performed for mouse (KPC) and human patient-derived PDAC organoids as indicated. (A) Key features of cell polarity were observed such as luminal microvilli, tight junctions and adherens junctions. The position of the lumen is denoted (**L**) along with the approximate positions of cell-cell junctions (▲▲) and the basal cell surface in proximity to the ECM (**$**). (C) Single arrowheads (▲) indicate the approximate position of some hemidesmosomes (indicated from the extracellular side) which are located at the plasma membrane (*), and often positioned proximal to a layer of basement membrane. (C) TEM was performed for normal human pancreas. The left-hand panel shows a lower magnification overview of a pancreatic ductal structure with lumen (L) and two cell nuclei (N). Ductal structures are surrounded by a basement membrane (**$**). The right-hand panel illustrates hemidesmosomes (▲) positioned at the basal surface of ductal cells in close proximity to a prominent basement membrane (**$**).

Another feature of the PDAC cell models highlighted by electron microscopy was the presence of desmosomes. In the HPDE cell line H6c7, in the absence or presence of KRas G12V, desmosomes were abundant, both in 2D and 3D culture models (Fig S2A and B). Furthermore, desmosomes were also detected in human patient-derived organoids and normal human pancreas (Fig S2C and D). In contrast, desmosomes were not detected in mouse organoids derived from the KPC mouse. These findings support previous reports of desmosomes in the human pancreas [56–58] and other human PDAC cell lines [59,60]. As desmosomes are known to play an important role in cancer progression [61,62], these results indicate differences between the cell-cell adhesion systems used in human PDAC and the KPC mouse model.

### Pancreatic cancer SUIT-2 cells form hemidesmosomes in 2D culture

To assess integrin α6β4 expression and hemidesmosome formation further, we tested integrin β subunit expression by flow cytometry using four commonly-used human PDAC cancer cell lines, SUIT-2, Panc1, MiaPaCa2 and KP4 [63,64] (Fig S3). Expression of cell surface integrin β1 and β5 was detected in all cell lines. In contrast, integrin β3 expression was very low or absent. Moreover, integrin β4 exhibited variable expression, with SUIT-2 the only cell line to demonstrate significant integrin β4 expression (Fig S3), which was confirmed by western blotting (Fig 7A).

**Figure 7:**
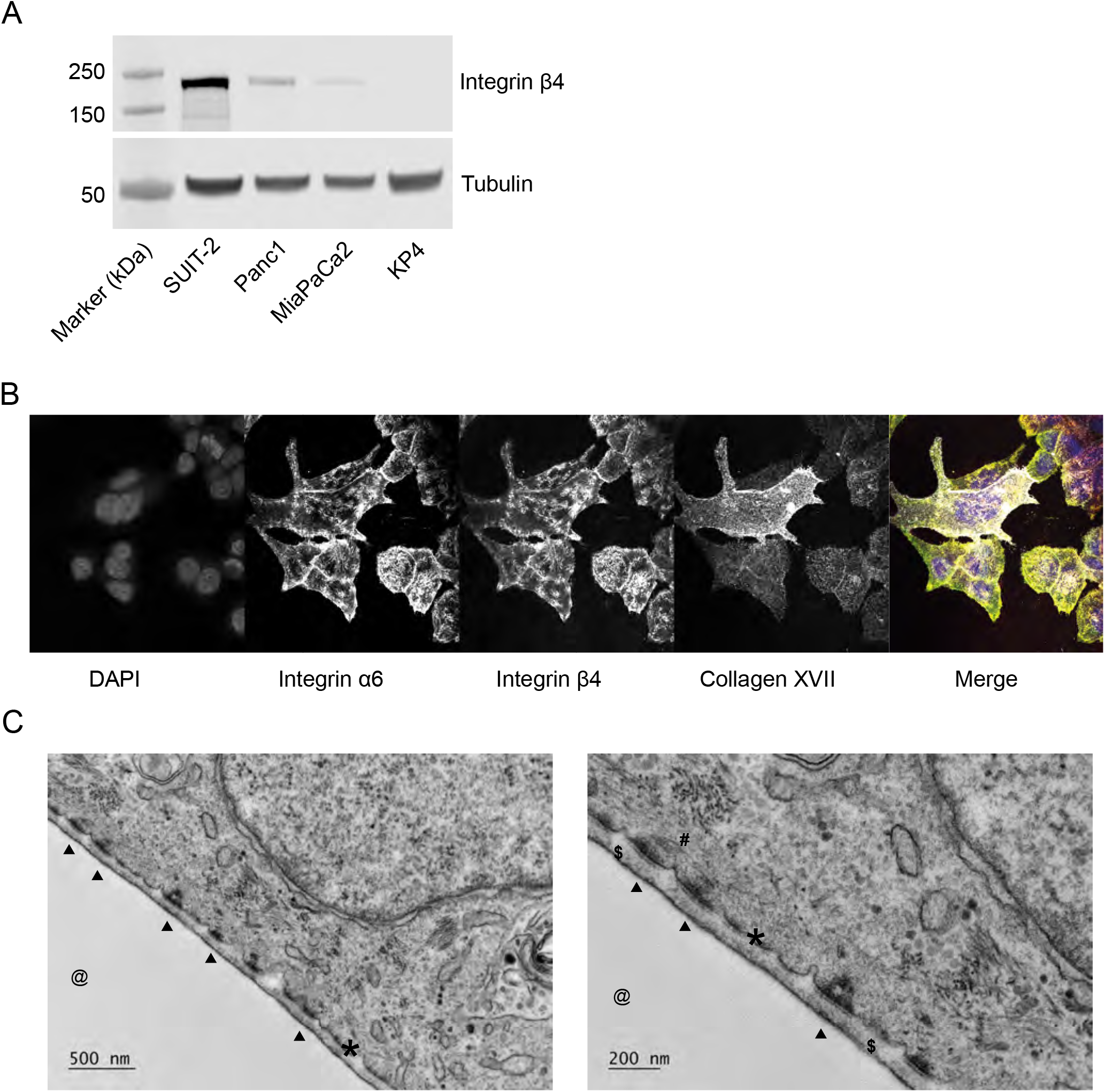
The human PDAC cell line SUIT-2 expresses integrin β4 and forms hemidesmosomes in 2D culture. (A) The expression of integrin β4 was assessed in SUIT-2, Panc1, MiaPaCa2 and KP4 PDAC cells by western blotting. Tubulin was used as a loading control. (B) SUIT2 cells were cultured for 7 days on glass coverslips and immunofluorescence imaging performed cells as indicated. (C) SUIT-2 cells were cultured on Aclar for up to 7 days. Transverse sections of the SUIT-2-ECM interface were prepared and imaged by TEM. The right-hand image shows a higher magnification of the same area. Cells formed a flattened basal surface with a thin layer of ECM (**$**) proximal to the area where the Aclar film (**@**) would have occupied. Arrowheads (▲) indicate the approximate position of some hemidesmosomes (indicated from the extracellular side) which are located at the plasma membrane (*) and link to cytoplasmic cytokeratin filaments (**#**). Images are orientated with the cell-ECM interface towards the bottom left.

To test for the formation of hemidesmosomes in the integrin β4-expressing cell line, immunofluorescence imaging was performed, and transverse sections of the SUIT-2-ECM interface were prepared and imaged by TEM. These analyses revealed a typical hemidesmosome-like localisation of integrin β4 (Fig 7B), and electron-dense hemidesmosome structures at the basal ECM interface of SUIT-2 cells, which were linked directly to prominent cytokeratin filaments (Fig 7C). Thus, ultrastructural analysis demonstrated that the integrin β4-expressing cell line, SUIT-2, also form hemidesmosomes in 2D culture and therefore expand the relevance of the observations of hemidesmosome formation to other PDAC cell models.

### HPDE and pancreatic cancer SUIT-2 cells require integrin β4 for proliferation

To assess the relevance of hemidesmosome formation and integrin β4 expression to HPDE and PDAC cell function, integrin β4 expression was depleted in HPDE and SUIT-2 cells using an siRNA knockdown approach. SUIT-2 and HPDE cells were used as they express integrin β4 and form hemidesmosomes. An almost complete depletion of integrin β4 was achieved in both HPDE and SUIT-2 cells (Fig S4). Cell proliferation was significantly reduced in both SUIT-2 and HPDE integrin β4 siRNA transfected cells compared to those transfected with control siRNA (Fig 8A and B). Furthermore, the number of SUIT-2 cells in S-phase, as determined by incorporation of EdU into replicating DNA, was significantly reduced by integrin β4 knockdown (Fig 8C). As this was indicative of changes in cell cycle progression, the proportion of SUIT-2 cells in G1, S or G2 was determined by flow cytometry. Knockdown of integrin β4 resulted in a significant reduction of SUIT-2 cells in S-phase, alongside a significant increase of cells in G1, demonstrating that depletion of integrin β4 levels induces G1 arrest in pancreatic epithelial cells (Fig 8D). These data demonstrate the functional significance of integrin β4 expression in HPDE and PDAC cells and suggest a role for hemidesmosomes in integrin β4-induced signal propagation to control cell cycle progression.

**Figure 8:**
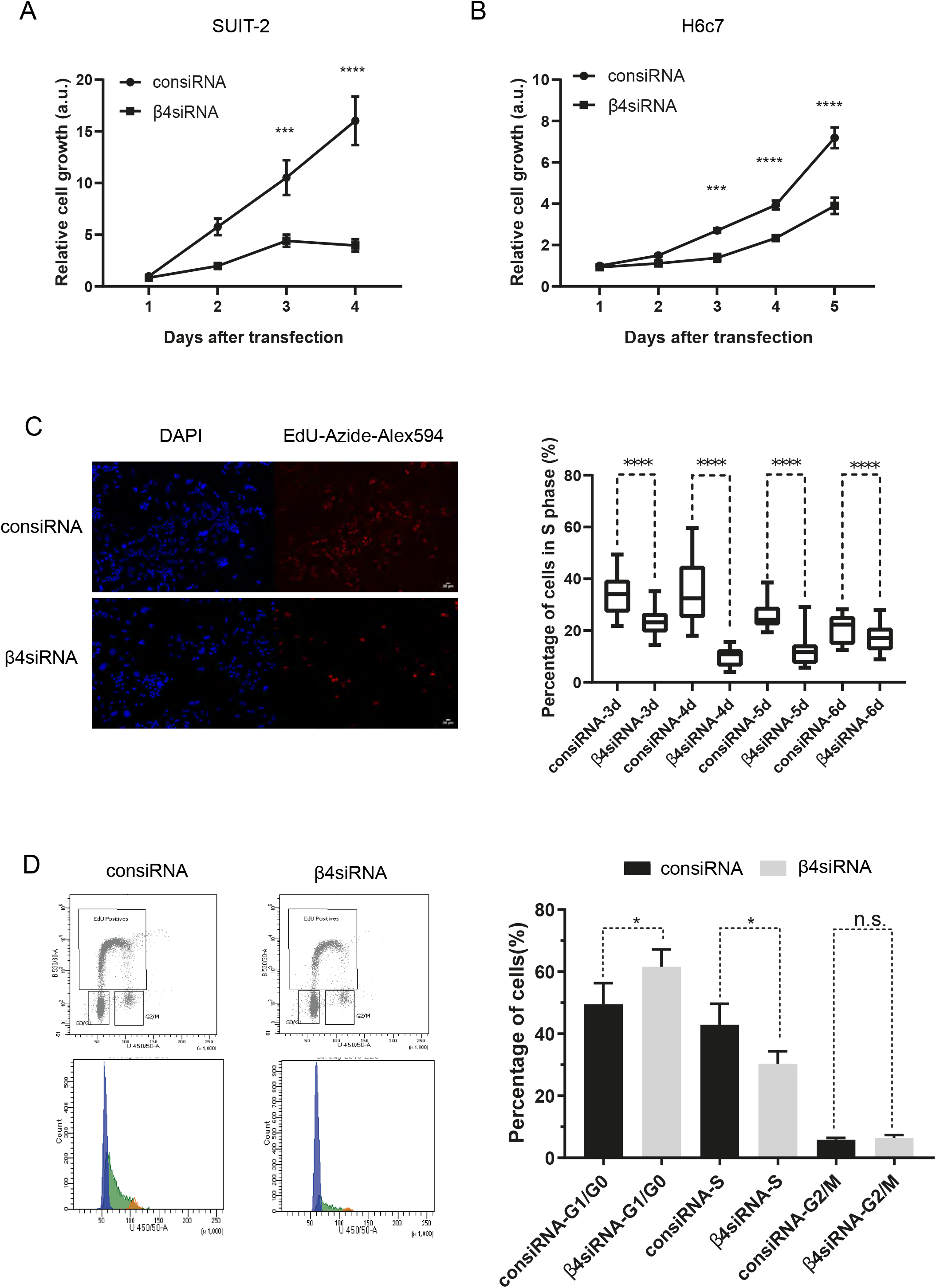
Depletion of integrin β4 reduces HDPE and SUIT-2 cell proliferation. (A) and (B) SUIT-2 cell proliferation was assessed over 5 days in culture after siRNA mediated depletion of integrin β4 (β4siRNA) compared to control siRNA (consiRNA). (C) EdU incorporation in SUIT-2 cells was assessed following siRNA-mediated depletion of integrin β4 and the percentage of cells in S phase calculated from three to six days after knockdown. (D) Cell cycle analysis by was performed using flow cytometry in SUIT-2 cells following siRNA mediated depletion of integrin β4. The proportion of cells in G1/G0, S and G2/M were calculated.

## Discussion

In this study, we analysed the integrin-based adhesion systems used by a variety of PDAC cell models and our major findings are: (1) HPDE cells primarily employ integrin α6β4 to adhere to their ECM; (2) α6β4 is assembled into hemidesmosomes in culture that are remarkably complete in composition and structure; and (3) disruption of hemidesmosomes by knockdown of β4 blocks the proliferation of HPDE cells. These findings suggest that the hemidesmosome adhesome may be exploited as a therapeutic target for PDAC and that HPDE cells provide an excellent model system for the study of both hemidesmosome assembly and function.

Using proteomics, we defined the ECM produced by HPDE cells, which contained abundant basement membrane components such as laminin 332. We also defined the adhesome of HPDE cells and demonstrated they form hemidesmosomes in 2D and 3D culture systems. No significant changes in hemidesmosome components were observed upon expression of mutagenic KRas G12V. We also demonstrated the formation of hemidesmosomes in another human PDAC cell line (SUIT-2), and in both mouse and human PDAC organoids. Furthermore, we showed that integrin β4 was critical for HPDE and SUIT-2 cell proliferation.

These data highlight the importance of laminins and laminin-binding adhesion mechanisms in a variety of PDAC models, and complements our recent study that defined a synthetic 3D model for the propagation of pancreatic ductal adenocarcinoma organoids [65]. The role of laminins in PDAC has been under-appreciated in the context of the well-characterised abundance of stromal collagens [4,6,66]. Our recent study defined the ECM changes that occur through PDAC progression, to inform the ECM adhesive cues provided in the synthetic 3D hydrogel scaffold, which highlighted the role of laminins. Importantly, we demonstrated a key role for laminin-cell interactions in the growth of PDAC-derived organoids and laminin-332 was upregulated in both human and murine PDA, which correlated with patient outcome [65]. In addition, we demonstrated that mouse and human pancreatic cancer cells attached and spread to laminins via integrins α3β1 and α6β1 [65]. As the use of organoids has been proposed as important for the future direction of PDAC investigations [55,67], the discovery of the formation of hemidesmosomes and the prominent role of laminin-cell interactions in these systems, and a variety of other PDAC cell models, is important.

Numerous studies have demonstrated important roles for laminins, laminin-binding integrins and hemidesmosome components such as collagen XVII in cancer [68–70]. The expression of laminin-332 and integrin β4 has also been reported in PDAC, and hemidesmosome formation has been reported in pancreatic exocrine tissue [71–75]. In this present study, we build and extend this knowledge by showing at the ultrastructural level that hemidesmosomes form in 2D and 3D for HPDE cells, and also murine and human PDAC organoids. The fact that we did not observe differences in the abundance of the hemidesmosomal components upon expression of mutagenic KRas G12V is consistent with the limited oncogenic potential originally reported for this cell line [41] and that additional mutations may be required to induced a complete malignant transformation [76]. This indicates that the early KRas G12V mutation in PDAC likely does not act to modulate adhesion signalling via altered hemidesmosome formation.

Finally, we show here that H6c7 HPDE cells are a good model cell line for formation of hemidesmosomes in 2D and 3D culture. All of the major hemidesmosome components were expressed and assembled into a fully developed plaque structure with intermediate filament attachment intracellularly, as well as all the recognised hemidesmosomes-associated structures in the ECM. This contrasts with a previous report that stated cells in culture do not form hemidesmosomes [44]. In fact, the literature reports several cell types, including rat bladder cancer (804G), human squamous cell carcinoma (DJM-1), mouse gingival epithelial (GE1) and mouse mammary tumour (RAC-11P/SD) cell lines, as forming hemidesmosomes at the ultrastructural level in 2D culture [77–81]. In addition, human breast MCF10A cells formed hemidesmosomes as acini in 3D culture [82]. Despite the expanding study of integrin adhesomes via IAC isolation and proteomics [83,84], only one study to date has reported the identification of integrin α6β4 from keratinocytes and two human oral squamous cancer cell lines [85]. Whilst this study reported the detection of some hemidesmosomal components such as collagen XVII and plectin from keratinocytes, it did not capture the full hemidesmosome repertoire, or claim to have isolated hemidesmosomes. We therefore propose that the dataset from this study is the first proteomics-based hemidesmosome adhesome to be reported. Interestingly, we identified FAT1 as an abundant non-hemidesmosome component in HPDE IACs in agreement with Todorovic et al 2010 [85]. FAT1 has recently been described as a having a role in tumour progression in squamous cell carcinoma through regulation of a hybrid epithelial-to-mesenchymal transition state [86]. This highlights the relevance of other ventral membrane proteins identified in these HPDE IAC datasets that may have a relevance to PDAC or other cancers.

In summary, using proteomics, we defined the ECM produced by HPDE cells, which contained abundant basement membrane components such as laminin-332. We also defined the adhesome of HPDE cells and demonstrated they form hemidesmosomes in 2D and 3D culture systems. No significant changes in hemidesmosome components were observed upon expression of mutagenic KRas G12V. We also demonstrated the formation of hemidesmosomes in another human PDAC cell line (SUIT-2), and in both mouse and human PDAC organoids. Finally, we demonstrated a functional role for integrin β4 in the regulation of HPDE and SUIT-2 cell proliferation, highlighting the potential of developing therapeutic strategies that target hemidesmosome components in PDAC.

## Experimental Procedures

### Reagents

Primary antibodies used for immunofluorescence microscopy were mouse monoclonal anti-β4 (3E1, Millipore), mouse monoclonal anti-dystonin (Cat #55654, Abcam), rabbit anti-E-cadherin (24E10, Cell Signaling Technology), mouse monoclonal directed against collagen VII (LH7.2, Abcam) and rabbit monoclonal anti-collagen XVII (ab184996, Abcam). Secondary antibodies (anti-mouse IgG Alexa Flour 488 and anti-rabbit IgG Alexa Flour 488) were from Invitrogen.

Primary antibodies for immunoblotting (at 1:1000 dilution) were monoclonal mouse anti-β1 integrin (JB1A, Millipore), monoclonal rabbit anti-β4 integrin (D8P6C, Cell Signaling Technology), polyclonal rabbit anti-α6 integrin (Cat #3750, Cell Signaling Technology), monoclonal rabbit anti-Ras (D2C1, Cell Signaling Technology), mouse anti-tubulin (DM1A, Sigma), mouse anti-paxillin (349, BD Biosciences) and monoclonal rabbit anti-collagen XVII (ab184996, Abcam). Secondary antibodies were goat anti-mouse IgG conjugated to Alexa Fluor 680 and goat anti-rabbit IgG conjugated to Alexa Fluor 800 (Life Technologies). Actin filaments were visualised by Alexa Fluor 594-conjugated phalloidin (Invitrogen).

Primary monoclonal anti-integrin antibodies for flow cytometry were mouse anti-α1 (TS2/7, Abcam), mouse anti-α2 (JA218 [87]), mouse anti-α4 (HP2/1, Abcam), mouse anti-α5 (JBS5, Millipore), rat anti-α6 (GoH3, Abcam), mouse anti-αV (17E6, Abcam), mouse anti-β1 (TS2/16, Invitrogen), mouse anti-β4 (3E1, Millipore), mouse anti-αVβ3 (LM609, Millipore), mouse anti-αVβ5 (P1F6, Millipore), Mouse IgG (Sigma), rat IgG (Sigma), rabbit F(ab’)2 anti-mouse IgG conjugated to FITC (STAR9B, BioRad), and rabbit F(ab’)2 anti-rat IgG conjugated to FITC (STAR17B, BioRad).

Primary monoclonal anti-integrin antibodies for immunoprecipitation were rat anti-α6 (GoH3, Abcam), mouse anti-β1 (TS2/16, Invitrogen), mouse anti-β4 (3E1, Millipore), mouse IgG (Sigma), and rat IgG (Sigma).

### 2D and 3D cell culture

The H6c7 cell line was used as a normal HPDE cell model. HPV-immortalised, human H6c7-pBABE and H6c7-KRasG12V cell lines were provided by M.S. Tsao, Ontario Cancer Institute, Canada [40]. The retroviral vector pBabepuro-KRAS4BG12V contained the human KRAS4B oncogene (KRASþ) cDNA with a mutation in codon12 (GTT to GTT). Cells were maintained in a 5% (v/v) CO_2_ humidified atmosphere at 37°C, in Keratinocyte Basal Medium supplemented with BPE, EGF, insulin, hydrocortisone and GA-1000 (Lonza) or Keratinocyte-SFM supplemented with L-glutamine, EGF, BPE and antibiotic-antimycotic (Life Technologies).

HPDE cells for 3D culture were incorporated into Matrigel/alginate gels and grown as described [51,88]. Briefly, 12 mg/ml growth factor reduced Matrigel (cat # 354230, BD Biosciences) was combined with 25 mg/ml alginate (Pronova SLG 100) resuspended in Dulbecco’s-modified Eagle’s medium (DMEM) on ice at a 2:1 ratio. HPDE cells were detached with trypsin/EDTA and 1×10^5^ cells per gel mixed with Matrigel/alginate mixtures. 1.22M calcium sulphate (CaSO4.2H_2_O) slurry was added in 50 µl DMEM to achieve a final concentration of 2.4 mM or 24 mM to generate medium and stiff gels, respectively. Rapid mixing of CaSO_4_ with Matrigel/alginate gels was achieved using 1 ml syringes connected via female-female luer lock couplers (Sigma, Superlco 21015). Mixed gels were immediately dispensed into wells of a 24-well plate, pre-coated with 50 µl Matrigel, and allowed to set for 30 minutes at 37°C before addition of HPDE growth medium.

SUIT-2, Panc1, MiaPaCa2 and KP4 cells [63,64,89] were obtained from ATCC and maintained in DMEM supplemented with 10% (v/v) fetal bovine serum (FBS; Life Technologies, Carlsbad, CA) and 2 mM L-glutamine. Cells were maintained at 37°C in a humidified atmosphere with 5% (v/v) CO2.

### Mouse and human organoid culture

Organoid growth and propagation conditions were as described by Below et al. [65]. In brief, murine pancreatic organoids (mPDOs) were isolated from tumour-bearing KPC mice from minced and enzymatically-digested tumour tissue. Human pancreatic organoids (hPDOs) were established from ultrasound-guided biopsy (EUS) or resected pancreatic cancer specimens. Cells were seeded in Matrigel and for passaging, mPDOs and hPDOs were separated from Matrigel by mechanical dissociation and seeded in a 1:6 (mPDO) or 1:2-1:4 (hPDO) split ratio into Matrigel droplets.

Patient research samples were obtained from the Manchester Cancer Research Centre (MCRC) Biobank with informed patient consent (www.mcrc.manchester.ac.uk/Biobank/Ethics-and-Licensing). The MCRC Biobank is licensed by the Human Tissue Authority (license number: 30004) and is ethically approved as a research tissue bank by the South Manchester Research Ethics Committee (Ref: 07/H1003/161+5).

### Ras activation assay

Assays were performed using Active Ras Detection Kit (Cell Signalling Technology), as per manufacturer’s guidelines. Briefly, cells were washed with Dulbecco’s phosphate-buffered saline (PBS), and lysed with the addition of complete protease inhibitor (Roche). Samples were centrifuged at 22,000 x g for 10 minutes at 4°C, and the supernatant applied to spin cups containing GST-Raf1-RBD or GST-control beads for 1 hr at 4°C. Samples were centrifuged at 6000 x g, washed three times with lysis buffer and eluted by addition of 2x SDS reducing sample buffer (50 mM Tris-HCl, pH 6.8, 10% (w/v) glycerol, 4% (w/v) sodium dodecylsulfate (SDS), 0.004% (w/v) bromophenol blue, 8% (v/v) β-mercaptoethanol). Samples were resolved by SDS-PAGE and immunoblotted as indicated.

### IAC isolation

IACs were isolated as described previously [42] without the use of the DTBP (Wang and Richard’s Reagent) protein crosslinker. Briefly, HPDE cells were cultured in full medium on two 10 cm diameter tissue culture dishes per condition for 7 days. Cells were washed twice with PBS and cell bodies removed by a 1 minute incubation with extraction buffer (50 mM Tris-HCl, pH 7.6, 150 mM NaCl, 0.5% (w/v) SDS, 1% (v/v) Triton X-100, 1% (w/v) sodium deoxycholate, 5 mM EDTA). Denuded cells were subjected to high-pressure water wash (30 seconds) to removed nuclei. IACs bound to the substrate were recovered in adhesion recovery solution (125 mM Tris-HCl, pH 6.8, 1% (w/v) SDS, 150mM dithiothreitol) and mixed with an appropriate volume of reducing sample buffer at 70 °C for 10 minutes before SDS-PAGE and immunoblotting or visualised with InstantBlue Coomassie stain (Expedeon) and processed for mass spectrometry (MS) as described below.

### Secreted protein collection

Serum-free growth medium, incubated for 72 hrs with confluent cells on 10 cm diameter tissue culture dishes was collected, passed through a 0.45 µm syringe filter to remove cells, and concentrated 40x by passing through VIVASPIN 20 MWCO 10 kDa centrifugal concentrator columns (Sartorius Stedim Biotech), as per manufacturer’s guidelines. The concentrated samples were resolved by SDS-PAGE, InstantBlue Coomassie stained and processed for MS as described below.

### CDM isolation

CDMs were generated as previously described [90–92]. In brief, HPDE cells were cultured from ∼50% confluency for 7 days, on 10 cm diameter tissue-culture dishes, washed in PBS, and lysed in extraction buffer (20mM NH4OH, 0.5% Triton X-100, in PBS) for 2 min at room temperature. CDMs were incubated with 10µg/ml DNase I at 37°C for 30 min, and proteins recovered in 2x reducing SDS buffer by scraping. Samples were resolved by SDS-PAGE, InstantBlue Coomassie stained and processed for MS as described below.

### MS sample preparation

Proteins were subjected to SDS-PAGE using 4-12% Bis-Tris gels (Life Technologies) for 3 minutes at 200 V, stained with InstantBlue Coomassie stain, and washed with distilled H_2_O overnight at 4°C. The gel band containing protein was excised and subjected to in-gel tryptic digestion in a perforated 96-well plate, as previously described [22]. Peptides were desalted using 1 mg POROS Oligo R3 beads (Thermo Fisher) as described [93], prior to MS analysis.

### MS data acquisition

Peptide samples were analysed by LC-MS/MS using an UltiMate 3000 Rapid Separation LC system (Thermo Fisher Scientific) coupled online to an Orbitrap Elite mass spectrometer (Thermo Fisher Scientific). Peptides were concentrated and desalted on a Symmetry C18 preparative column (20 mm × 180 μm 5-μm particle size, Waters) and separated on a bridged ethyl hybrid C18 analytical column (250 mm × 75 μm 1.7-μm particle size, Waters) using a 45-min linear gradient from 1% to 25% or 8% to 33% (v/v) acetonitrile in 0.1% (v/v) formic acid at a flow rate of 200 nl min^−1^. Peptides were selected for fragmentation automatically by data-dependent analysis.

### MS data analysis

Tandem mass spectra were extracted using extract_msn (Thermo Fisher Scientific) or ProteoWizard [94] executed in Mascot Daemon (version 2.5.1; Matrix Science). Peak list files were searched against a modified version of the Uniprot human database, using Mascot (version 2.5.1; Matrix Science) [95]. For secreted protein and CDM analyses, Uniprot human database (release-2018_01) was used. For HPDE IAC analysis Uniprot human database (release-2016_04) was used. Carbamidomethylation of cysteine was set as a fixed modification; oxidation of methionine was allowed as a variable modification. For secreted protein and CDM analyses, hydroxylation of proline and lysine were allowed as additional variable modifications. Only tryptic peptides were considered, with up to one missed cleavage permitted. Monoisotopic precursor mass values were used, and only doubly and triply charged precursor ions were considered. Mass tolerances for precursor and fragment ions were 5 ppm Da and 0.5 Da, respectively. MS datasets were validated using statistical algorithms at both the peptide and protein level implemented in Scaffold (version 4.4.7, Proteome Software) [96,97]. For the IAC dataset protein identifications were accepted upon assignment of at least two unique validated peptides with ≥90% probability, resulting in ≥99% probability at the protein level. These acceptance criteria resulted in an estimated protein false discovery rate of <0.01% with zero decoys. For the CDM and secreted protein datasets protein identifications were accepted upon assignment of at least two unique validated peptides with ≥95% probability, resulting in ≥99% probability at the protein level. These acceptance criteria resulted in an estimated protein false discovery rate of <0.01% with zero decoys, and 0.02% with one decoy for CDM and secreted protein datasets respectively. Comparison of datasets with the IAC adhesome and matrisome were performed using the tools reported previously [43] and included ECM components of the consensus adhesome.

### MS data quantification

For the IAC MS dataset, relative protein abundance was calculated using peptide intensity using Progenesis LC-MS (Non Linear Dynamics) with automatic alignment as previously described [93]. Orbitrap MS raw data was imported into Progenesis LC-MS to acquire intensity data. Features with number isotopes >2 and charge states of <5 were used to filter for peptide identifications and exported to an in-house Mascot server as described above. Mascot results were imported to Scaffold as above and peptide and protein identification thresholds were set to 90% and 99% confidence, respectively. Data were exported from Scaffold as a spectrum report, and imported into Progenesis LC-MS to assign peptide identifications to features. Protein identifications with quantification and assigned statistical q-values were then exported to be analysed. For secreted protein and CDM analysis, protein abundance was calculated as spectral counts as reported by Scaffold analysis and statistical comparisons made using QSpec [98]. Data were visualised as volcano plots using the online version of VolcaNoseR (https://huygens.science.uva.nl/VolcaNoseR) [99].

### MS data deposition

The mass spectrometry proteomics data have been deposited to the ProteomeXchange Consortium via the PRIDE partner repository with the dataset identifiers PXD027803 (Project DOI: 10.6019/PXD027803), PXD027823 (Project DOI: 10.6019/PXD027823) and PXD027827 (Project DOI: 10.6019/PXD027827) [100].

### Gene ontology (GO) enrichment analysis

Official gene symbols were mapped to all protein identifications, and datasets were analysed using the online bioinformatic tools available via the Database for Annotation, Visualization and Integrated Discovery (DAVID; http://david.abcc.ncifcrf.gov/home.jsp) [101]. Only terms with enrichment value ≥1.5, Bonferroni-corrected P-value <0.05, EASE score (modified Fisher Exact P-value) <0.05 and at least two genes per term were considered.

### Immunoprecipitation

Cells were harvested and lysed for 10 minutes at 4°C with Pierce IP Lysis Buffer (ThermoFisher Scientific) supplemented with complete protease inhibitors (Roche). Cell debris was pelleted by centrifugation at 22,000 x g for 10 minutes at 4°C, and the supernatant incubated with Protein-G Sepharose for 1 hr at 4°C. Antibody/Protein-G Sepharose mixes were added for 30 minutes at room temperature, centrifuged at 2680 x g for 2 minutes at 4°C, and the supernatant discarded. Samples were then washed with lysis buffer before elution of bound proteins with 2x reducing SDS buffer for 10 minutes at 70°C. Samples were centrifuged, the supernatant collected, resolved by SDS-PAGE and immunoblotted.

### Immunoblotting

Unless otherwise specified, cells were lysed in 150 mM NaCl, 25 mM Tris-HCl, pH 7.4, 1 mM EDTA, 1% (v/v) NP-40, 5% (v/v) glycerol, 50 µg/ml leupeptin, 50 µg/ml aprotinin, 1 mM 4-(2-aminoethyl)-benzenesulfonyl fluoride (AEBSF) and 1x PhosSTOP phosphatase inhibitor cocktail (Sigma-Aldrich), and centrifuged at 10,000 x g for 10 minutes at 4 °C. Cell lysates were separated by SDS–PAGE (4-12% Bis-Tris gels, Thermo Fisher) under reducing conditions and transferred to nitrocellulose membrane (Whatman). Membranes were blocked for 60 minutes at room temperature using either casein blocking buffer (Sigma-Aldrich) or 5% (w/v) bovine serum albumin in 10 mM Tris-HCl, pH 7.4, 150 mM NaCl containing 0.05% (w/v) Tween-20 (TBST). Primary antibodies, diluted in blocking buffer or 5% (w/v) bovine serum albumin/TBST, were probed overnight at 4 °C and membranes washed using TBST for 30 minutes at room temperature. Secondary antibodies diluted in blocking buffer or 5% (w/v) bovine serum albumin/TBST were then incubated for 45 minutes at room temperature in the dark and membranes washed in the dark using TBST for 30 minutes at room temperature. Bound antibodies were visualised using an Odyssey infrared imaging system (LI-COR) and band intensities analysed using Odyssey software (LI-COR).

### siRNA knockdown of integrin β4

HPDE and SUIT-2 cells were transfected with siRNAs by using oligofectamine (Sigma-Aldrich) according to the manufacturer’s instructions. Knockdown of integrin β4 was performed by using SMARTpool reagents (L-008011-00-0005, Horizon) and ON-TARGETplus nontargeting siRNA (Horizon) was used as a negative control.

### Immunofluorescence and image analysis

Cells were cultured for up to 7 days on glass coverslips and fixed with 4% (w/v) paraformaldehyde for 15 minutes at room temperature and permeabilised with 0.2% (w/v) Triton X-100 in PBS for 20 minutes at room temperature. Coverslips were incubated with primary antibodies directed against proteins indicated in 2% (w/v) bovine serum albumin in PBS for 1 hour at room temperature. Cells were then incubated with fluorophore-conjugated secondary antibodies for 45 minutes at room temperature, stained with 1 μg/ml DAPI for 1 minute before washing and mounting onto glass slides. Images were acquired using an Olympus BX51 upright microscope with a 60x/0.65-1.25 UPlanFLN or 10x/0.30 UPlanFLN objective and captured using a Coolsnap EZ camera (Photometrics) through MetaVue software (Molecular Devices). Alternatively, images were acquired on an inverted confocal microscope (TCS SP5 Acousto-Optical Beam Splitter; Leica) by using a 63x objective (HCX Plan Apochromat, NA 1.25) and Leica Confocal Software (Leica). Image analysis was performed using ImageJ [102].

### EdU incorporation, cell proliferation and cell cycle analysis

To assess the proportion of proliferating cells, HPDE or SUIT-2 cells were either plated onto glass coverslips or tissue culture dishes and transfected as indicated with siRNA (Dharmacon on-target smartpool, Horizon Discovery). After 48 hours, cells were pulse-labelled with 10 µM EdU for 50 minutes, fixed and EdU-labelled using Click-it chemistry according to the manufacturer’s instructions (Thermo Fisher Scientific).

For imaging analysis, cells were counterstained with DAPI and phalloidin, washed three times with PBS containing 0.1% (w/v) Tween-20, and once with distilled H_2_O before mounting on coverslips by using ProLong diamond antifade reagent (Thermo Fisher) and imaging. Images were collected on a Zeiss Axioimager D2 upright microscope using a 10x/0.3 EC Plan-neofluar objective and captured using a Coolsnap HQ2 camera (Photometrics) through Micromanager software v1.4.23. The total number of DAPI-positive nuclei were counted and the proportion of these that were positive for EdU staining determined.

For analysis by flow cytometry, EdU-labelled cells were stained with FxCycle violet (Thermo Fisher) to label DNA content in the cell. Samples of 10,000 cells were then analysed using a BD LSR Fortessa flow cytometer and FlowJo to determine the proportion of cells in G1, S and G2.

### Flow cytometry

Cells were washed with PBS, detached with 1x trypsin-EDTA at 37°C and harvested by centrifugation at 280 x g for 4 min. Cell pellets were resuspended at 0.5-1 × 10^7^ cells/ml in PBS with 1% (v/v) fetal calf serum on ice. Primary antibodies were diluted in PBS plus 0.1% (w/v) sodium azide and incubated with cells at 10 µg/ml for 60 minutes at 4°C. Following two washes with PBS containing fetal calf serum and centrifugation at 280 x g for 4 min, cells were incubated with appropriate species-specific FITC-conjugated secondary antibodies for 30 minutes at 4°C. Cells were then washed three times with PBS containing fetal calf serum, centrifuged at 280 x g for 4 min, resuspended in PBS, fixed with 0.4% (v/v) formaldehyde in PBS and analysed on a Dako CYAN, or Beckman Coulter Cyan ADP FACS machine (Beckman Coulter).

### Electron microscopy

HPDE and SUIT-2 cells were grown on Aclar film (Agar Scientific) for seven days in culture medium and fixed with 4% (v/v) formaldehyde plus 2.5% (v/v) glutaraldehyde in 0.1M HEPES buffer (pH 7.2). Subsequently, samples were post-fixed with 1% (w/v) osmium tetroxide and 1.5% (w/v) potassium ferricyanide in 0.1M cacodylate buffer, pH 7.2 for 1 hour, then 1% (w/v) tannic acid in 0.1M cacodylate buffer, pH 7.2 for 1 hour and finally in 1% (w/v) uranyl acetate in distilled water for 1 hour. Samples were then dehydrated in an ethanol series infiltrated with TAAB Low Viscosity resin and polymerised for 24 hr at 60°C as thin layers on Alcar sheets. After polymerisation, Aclar sheets were peeled off and layers of polymerised resin with cells were re-embedded with the same resin as stacks. Sections were cut with a Reichert Ultracut ultramicrotome and observed with a FEI Tecnai 12 Biotwin microscope at 100kV accelerating voltage. Images were taken with a Gatan Orius SC1000 CCD camera.

### Human pancreas samples

Access to human pancreas samples for electron microscopy was obtained by following recommendations from the National Research Ethics Services (NRES). The protocol was ethically approved by the North-West Research Ethics Committee (Ref: 07/H1010/88).

## Abbreviations

CDM: cell-derived matrix
DTBP: dimethyl 3,3’-dithiobispropionimidate (Wang and Richard’s Reagent)
ECM: extracellular matrix
GO: gene ontology
HPDE: human pancreatic ductal epithelial
IAC: integrin adhesion complex
KPC: KrasG12D/WT; TP53R172H/WT; Pdx1-Cre mice
LC-MS/MS: liquid chromatography-tandem MS
MS: mass spectrometry
PDAC: pancreatic ductal adenocarcinoma

## Author contributions

Conceptualisation: JDH, JAA, CJ and MJH. Methodology: JDH, CJB, MCJ, AM and DK. Resources: AM, DK, DAO and MJD. Investigation: JDH, JZ, JB, MRC, JAA, CJB, AM, DAO, MJD, DRG and JDH. Writing: JDH and MJH (original draft); JDH, JZ, MRC, MCJ, AM, DRG, CJ and MJH (review & editing). Funding Acquisition: MJH and CJ. Supervision: JDH, MJH and CJ.

## Acknowledgments

This work was supported by a Cancer Research UK Programme Grant (C13329/A21671; to MJH and CJ), Cancer Research UK Institute Award (A19258; to CJ), Cancer Research UK Experimental Medicine Programme Award (A25236; to CJ), the Rosetrees Trust (M286; to CJ) and a European Research Council Consolidator Award (ERC-2017-COG 772577; to CJ). Megan Chastney was supported by a PhD studentship from BBSRC. The work was conducted within the Wellcome Centre for Cell-Matrix Research (core award 203128/Z/16/Z). The support of the Bio-MS mass spectrometry, flow cytometry, electron microscopy and bioimaging core facilities in the Faculty of Biology, Medicine and Health at the University of Manchester is gratefully acknowledged.

The mass spectrometers and microscopes used in this study were purchased with grants from BBSRC, Wellcome Trust and the University of Manchester Strategic Fund.

We thank E. Keevill (University of Manchester) for acquisition of MS data, J.N. Selley (University of Manchester) for bioinformatic support, P. March and R. Meadows (University of Manchester) for assistance with microscopy, X. Zhang and C. Hutton (Cancer Research UK Manchester Institute) for assistance with organoid culture. Monoclonal mouse antibody against human integrin β1 (clone JB1A) was kindly provided by J. A. Wilkins (University of Manitoba, Winnipeg, MB, Canada).

## Conflicts of interest

The authors declare no conflicts of interest.

## Appendix A. Supplementary data

**Supplementary Table 1** MS data - H6c7 HPDE IAC Scaffold and Progenesis QI Analysis

**Supplementary Table 2** MS data - H6c7 HPDE CDM Scaffold analysis

**Supplementary Table 3** MS data - H6c7 HPDE secreted protein Scaffold analysis

**Supplementary Table 4** HPDE CDM and secreted protein Gene Ontology analysis

---

**Figure S1:**
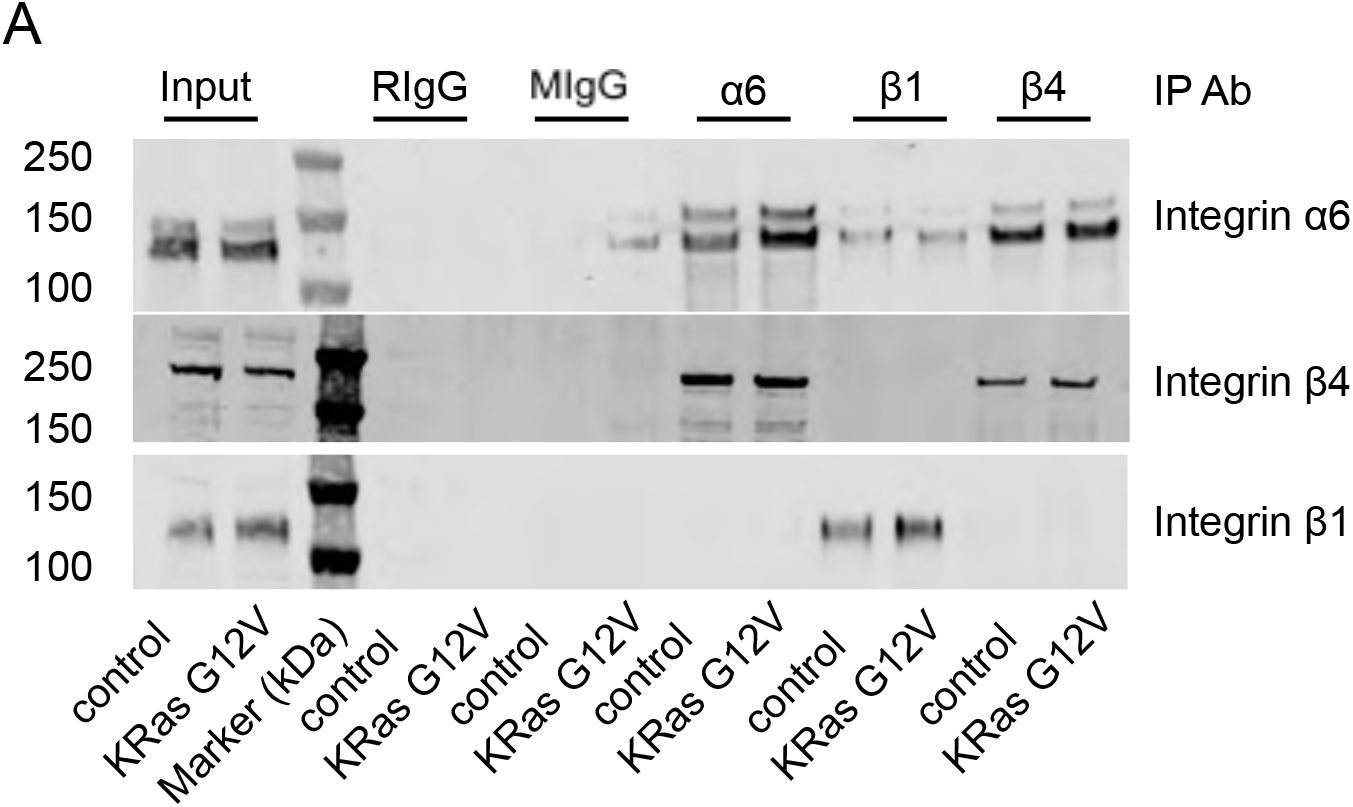
HPDE cells express the integrin α6β4 laminin-binding integrin subunits. Flow cytometry of integrin subunits (A) Immunoprecipitation was performed from wild-type (control) and mutant KRas-expressing (KRas G12V) HPDE lyates using anti-integrin α6, β1 and β4 antibodies, along with control rat (RIgG) and mouse (MIgG) polyclonal antibodies. SDS-PAGE and western blotting using anti integrin α6, β1 and β4 antibodies (under reducing conditions) confirmed the preferential association of α6 with the β4 subunit compared to β1. The sizes of molecular weight markers are indicated to the left of images.

**Figure S2:**
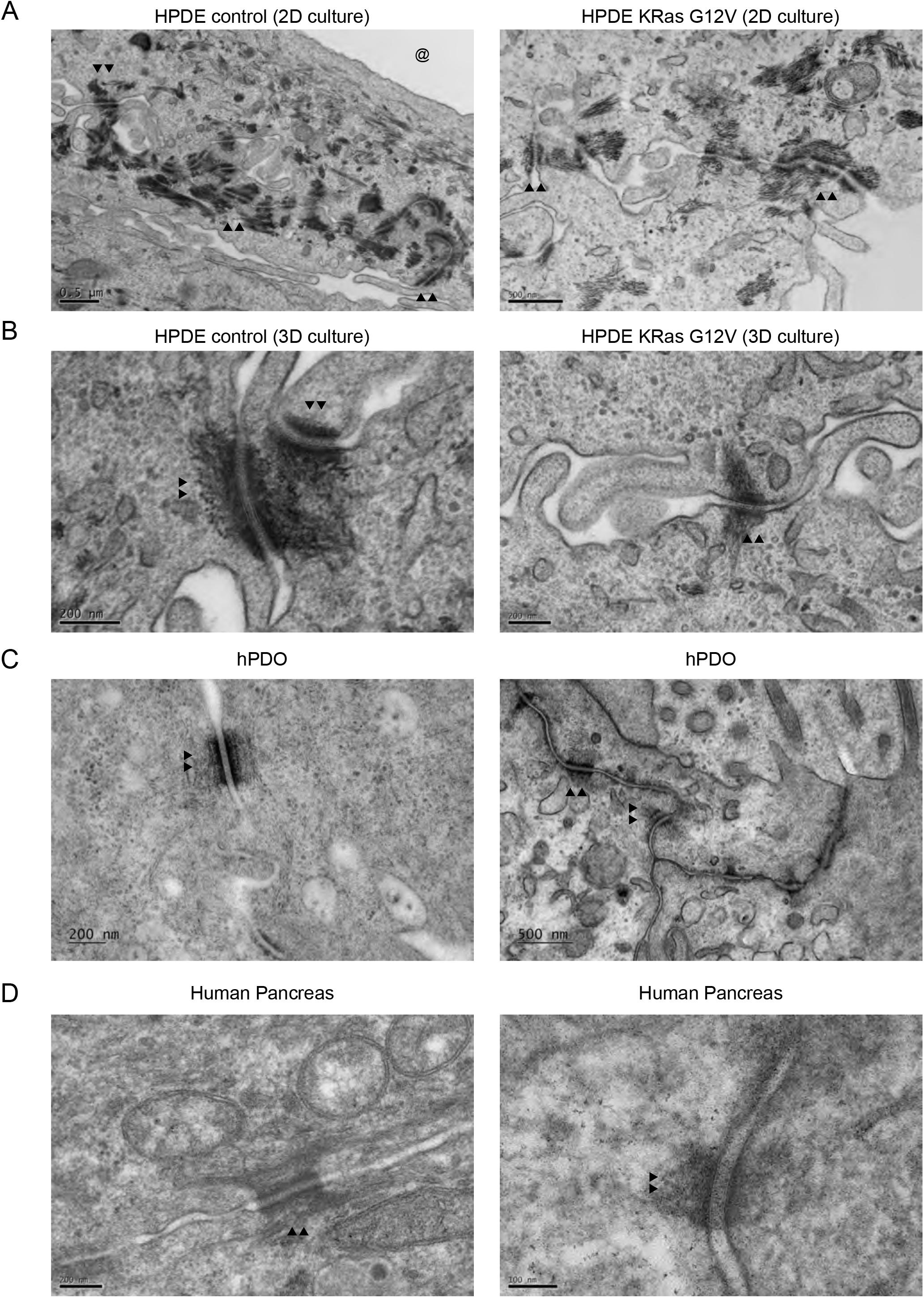
Desmosomes are observed in HPDE cells, human PDAC organoids and human pancreas. (A) Wild-type (control) and mutant KRas-expressing (KRas G12V) HPDE cells were cultured on Aclar for up to 7 days. Transverse sections of the HPDE-ECM interface were prepared and imaged by TEM. Cells formed flattened basal surface proximal to the area where the Aclar film (@) would have occupied. Double arrowheads (▲▲) indicate the approximate position of some desmosomes. (B) TEM was performed for wild-type (control) and mutant KRas-expressing (KRas G12V) HPDE cells after 60 hours in 3D culture (alginate / Matrigel). Double arrowheads (▲▲) indicate the approximate position of desmosomes. (C) TEM was performed for human patient-derived PDAC organoids. Double arrowheads (▲▲) indicate the approximate position of some desmosomes. (D) TEM was performed for normal human pancreas. Double arrowheads (▲▲) indicate the approximate position of some desmosomes.

**Figure S3:**
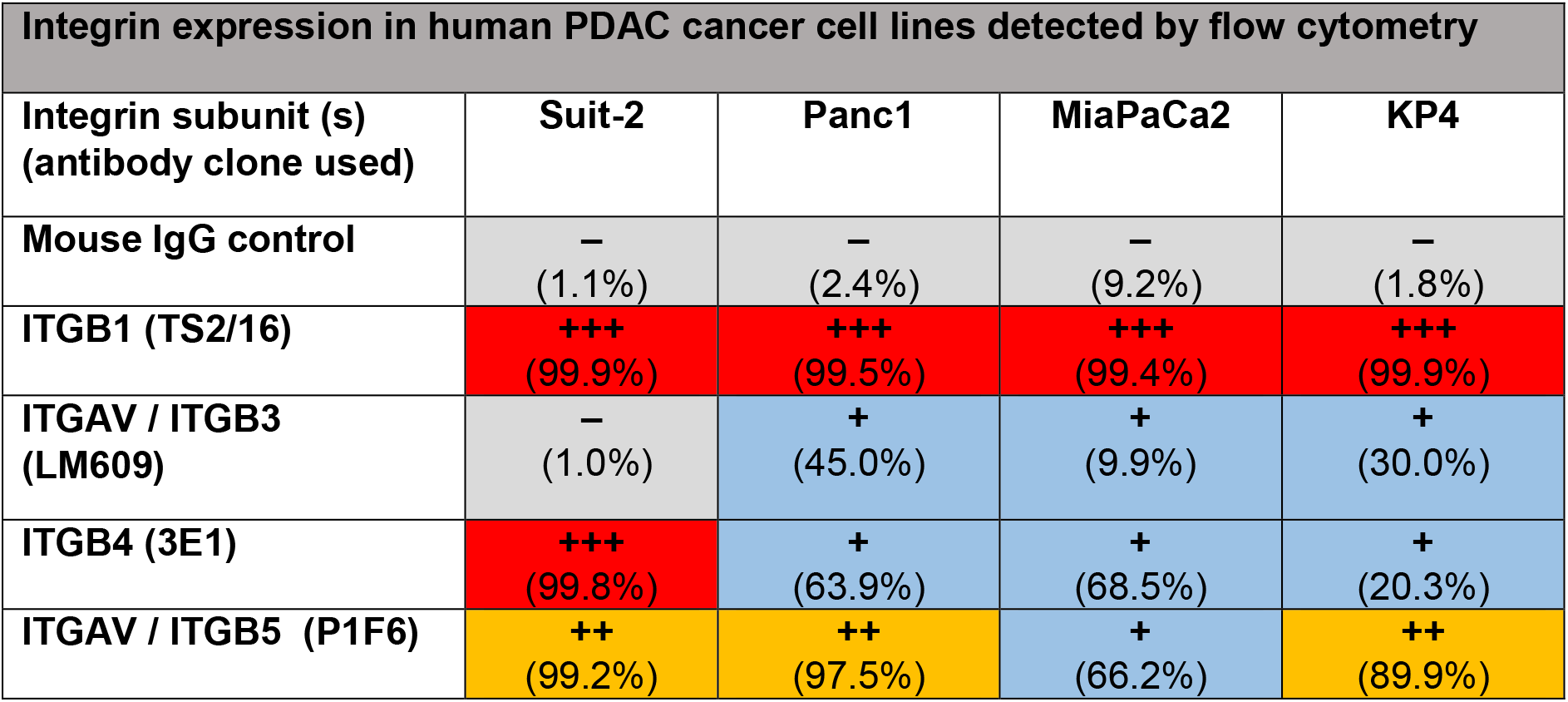
Integrin expression in human PDAC cancer cell lines as determined by flow cytometry. Flow cytometry of integrin subunits in SUIT-2, Panc1, MiaPaCa2 and KP4 cells. Values indicate integrin expression as median fluorescence intensities (+++ = 103-104; ++ = 103; + = 102-103; – = 102) with percent of cells expressing higher than MuIgG control in parenthesis. Values are indicative of two independent experiments.

**Figure S4:**
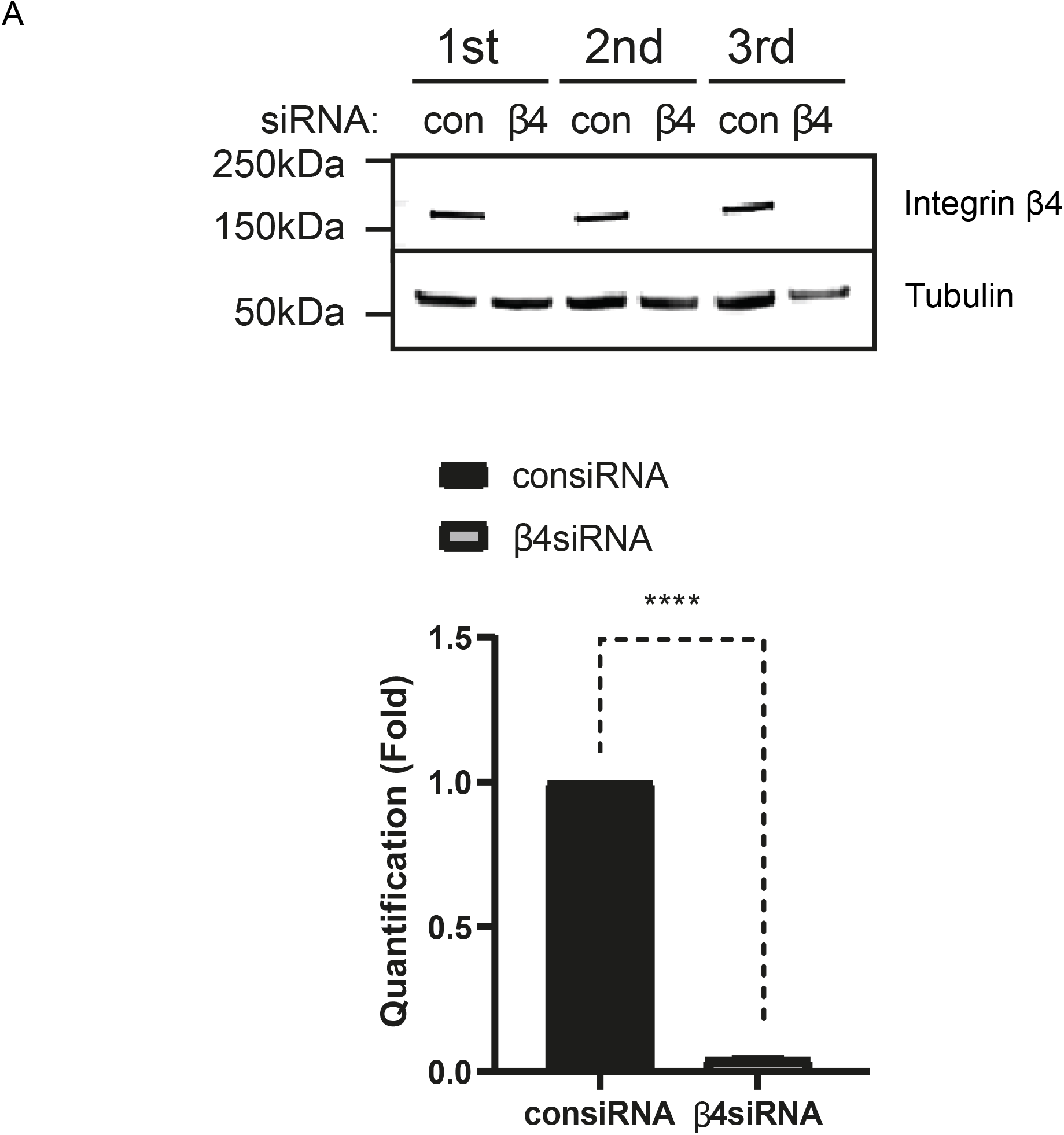
siRNA-mediated knockdown of integrin β4 in SUIT-2 cells. SUIT-2 cells were subjected to siRNA-mediated knockdown of integrin β4. Western blotting demonstrated reduced expression of integrin β4 siRNA compared to control siRNA (con). Tubulin was used as a loading control. Data from three independent repeats are shown and quantified.

## Notes

### Competing Interest Statement

The authors have declared no competing interest.

